# Modeling Dynamic Transcriptional Circuits with CRISPRi

**DOI:** 10.1101/225318

**Authors:** Samuel Clamons, Richard M. Murray

**Affiliations:** Caltech, Pasadena, CA, United States

## Abstract

Targeted transcriptional repression with catalytically inactive Cas9 (CRISPRi) promises to reproduce the functions of traditional synthetic transcriptional circuits, but with better orthogonality, programmability, and extensibility. However, CRISPRi lacks obvious cooperativity–a feature classically considered critical for several classic gene regulatory circuits. We use a simple dynamical model of CRISPRi to show that it can be used to build repressilators, toggle switches, and incoherent feed-forward loops. We also show that the function some of these circuits are expected to be sensitive to several key parameters, and we provide specifications for those parameters. Our modeling reveals key engineering requirements and considerations for the construction of dynamic CRISPRi circuits, and provides a roadmap for building those circuits.

## 1. Introduction

A central challenge of modern bioengineering is that of “programming” cells with complex, dynamic behaviors. Simple examples of genetically encoded dynamic functions include cell state oscillation [1, 2, 3], event detection and logging [4], molecular fold-change detection [5, 6], and signal level discrimination [7]. A common challenge when engineering complex behavior is the need for numerous *specific* interactions between components. In general, engineering specific, efficient, non-promiscuous reactions from scratch is difficult for a number of reasons, so bioengineers typically use natural systems whose components have built-in specificity and selectivity.

One such natural molecular system is that of the gene regulatory network. Synthetic gene regulatory networks exploit the ability of transcription factors to specifically control the actions of target promoters to “wire” together transcriptional units, much the same way microchip manufacturers use spatial arrangements to wire together silicon-based components like transistors and logic gates. Gene regulatory networks have been successfully used to build small circuits [1, 8, 9], but they have not been used to build systems with more than about a dozen regulators. Major barriers to scaling up genetic regulatory networks include a lack of orthogonal transcription factors (the largest verified-orthogonal library of repressors currently consists of about 16 genes [10]), mismatches in output and input levels between different regulators, and metabolic burden on the host cell [11].

One system that promises to scale better than classical genetic regulators is CRISPRi, a system of repression that uses a catalytically inactive mutant of the programmable endonuclease Cas9 (“dCas”). A dCas protein is inactive until loaded with a guide RNA (gRNA) containing a roughly 20-bp variable region. Once loaded, dCas will bind to any double-stranded DNA sequence matching the variable region of the gRNA, so long as it is immediately upstream of a short PAM region (NGG for the commonly-used *S. pyogenes* dCas9, but different for different dCas variants) [12]. Binding of dCas can interfere with prokaryotic transcription, either preventing initiation of transcription (if the gRNA is targeted within or immediately around a promoter) or blocking elongation (if the gRNA is targeted downstream of the promoter) [13].

CRISPRi repressors have several potential advantages over traditional transcription factors. The clearest advantage of CRISPRi is that it provides an almost limitless supply of orthogonal repressors. Another advantage of CRISPRi is the relative uniformity of CRISPRi repressors. Since many CRISPRi operators can be made using the same core promoter sequence, it might be expected that different CRISPRi repressors should act with similar input/output relationships.

Since it was first proposed [14], CRISPRi has been widely used for biophysical characterization of Cas9 [15, 16, 17, 18, 19, 20] and for control of host gene expression [21, 22, 23]. More rarely, CRISPRi has been used as a synthetic tool in eukaryotic systems. Layerable CRISPRi endpoint logic gates have been designed at least twice [10, 24], and circuits up to eight gates deep and utilizing up to a dozen gates in total have been constructed. CRISPRi has also been used to make circuits with simple dynamic behavior [25, 26] but has not yet been widely adapted to create scalable prokaryotic circuits.

We use a simple model to demonstrate that CRISPRi can be used to build both toggle switches and repressilators, despite the fact that CRISPRi displays no cooperativity. We predict that the CRISPRi repressilator should function under physiological conditions in *E. coli*, but only barely (an observation backed up by other experiments in the literature), and suggest several interventions that should make the CRISPRlator more robust.

## 2. A Model of CRISPRi

Most of our analyses will use a mass-action ODE model, even though there is good reason to believe that at least some components of a CRISPRi network will be present at low concentrations (< 10 molecules/cell), on the grounds that 1) ODE models are easier to write, simulate, understand, and analyze than stochastic models, and 2) ODE models can provide insight even in systems where the bulk assumptions of a mass-action may not be justified.

We describe dCas binding and unbinding explicitly rather than with a Hill approximation on the grounds that dCas binding and unbinding is relatively slow compared to typical timescales of biocircuit activity, so we cannot assume that dCas will come to equilibrium quickly.

### 2.1. A description of the model

A simple CRISPRi model, as shown in Figure 1, includes the following processes:

- Production of dCas;
- Production of gRNAs from free promoters;
- Leak production of gRNAs from dCas-bound promoters;
- Active degradation of free gRNAs (by RNAses);
- Global dilution;
- Maintenance of promoter copy number, and unbinding of gRNA:dCas complexes triggered by replication;
- Binding and unbinding of gRNAs from dCas;
- Binding and unbinding of gRNA:dCas complex from target promoters;

**Figure 1:**
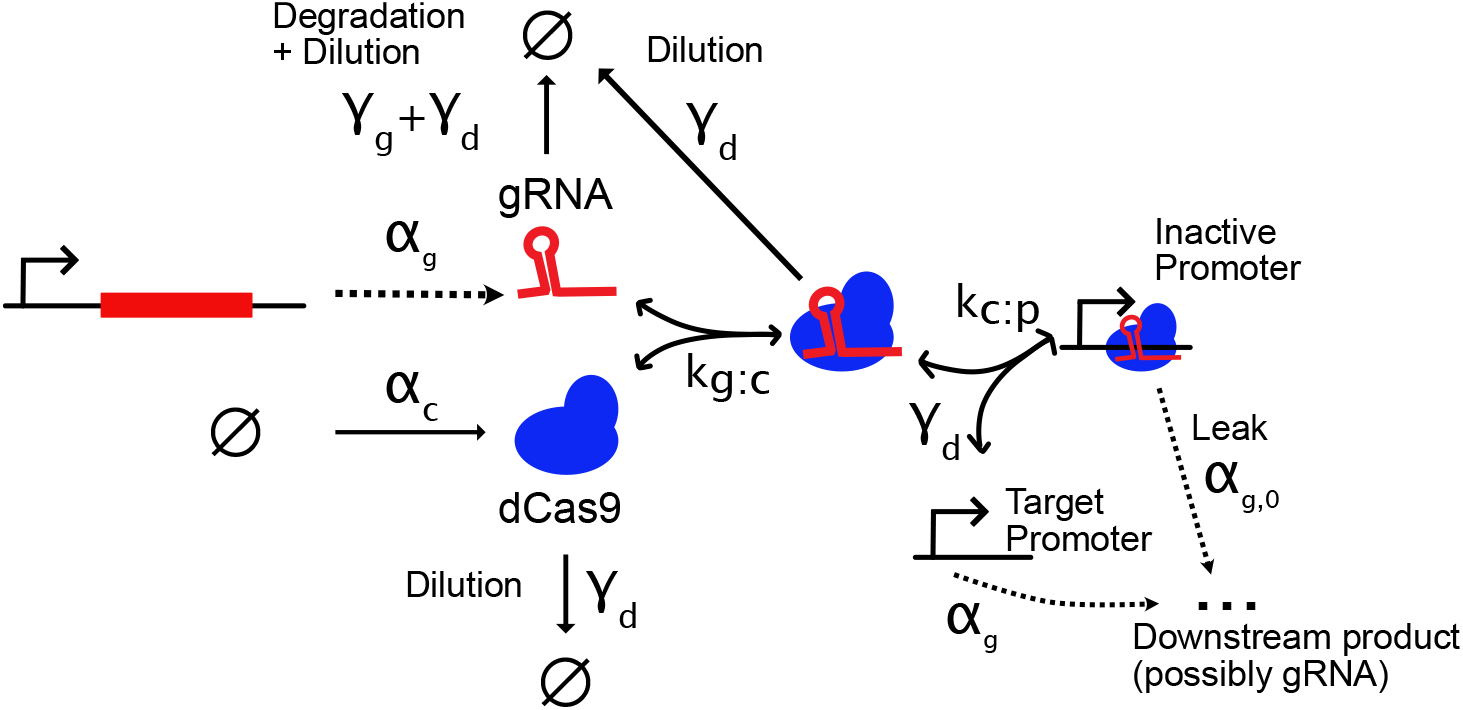
Diagrammatic representation of the CRISPRi model used in this report. Rates for active degradation of dCas and leak activity of dCas-bound promoters are set to zero in some simulations.

We take production of dCas to be constant; gRNA production is constant from unbound promoters, and constant from bound promoters with a lower rate (possibly zero). Guide RNAs (and, optionally, dCas and dCas complexes) are actively degraded at a rate proportional to their abundance. Binding reactions (and unbinding of gRNA from dCas) follow standard mass action binding and unbinding kinetics. Except where explicitly stated otherwise, all CRISPRi promoters are assumed to have identical dynamics (aside from the identity of their repressors).

Unbinding of dCas from its target promoters is a little unusual in our model. dCas binds extremely tightly to its targets. In bacterial cells, the rate of dCas unbinding from DNA is substantially slower than the rate of bacterial replication, even in non-lab-adapted strains with division times of over 100 minutes [20] (though possibly not [19]). This means that dCas effectively only unbinds as a consequence of DNA replication, with the bacterial replication machinery forcing dCas from its target. Any CRISPRi circuit with complex, non-monotonic dynamics will require either dilution, dCas degradation, or some mechanism that actively unbinds dCas from its DNA targets.

Accordingly, all models in this report apply global cell dilution to all components (including DNA, dCas, and complexes of the two), using a series of reactions of the form 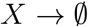, where *X* is any species in the model. A caveat is that total promoter concentrations need be held constant, as dilution of DNA is assumed to be equally balanced by replication. We implement this with a reaction *DNA_ij_* → *DNA_ij_* + *DNA_ij_* at the rate of dilution for each *DNA_ij_* in the model, where *DNA_ij_* is any unbound promoter for gRNA *i* that is repressed by gRNA *j*. DNA bound to dCas follows a combined replication/unbinding reaction *DNA_ij_*: *dCaS_j_* → *DNA_ij_* + *dCaS_j_*, again at the rate of dilution, where *dCaS_j_* is a dCas molecule complexed to gRNA *j*. This reaction represents the current understanding that dCas is removed from DNA by DNA replication.

Note that the reactions *DNA_ij_* → *DNA_ij_* + *DNA_ij_* and *DNA_ij_* → 0 make *DNA_ij_* only neutrally stable – they do not reject disturbances to DNA levels. The precarious balance this simple replication mechanism achieves is be sufficient for the deterministic simulations we performed, but would need to be modified if used for stochastic simulation [27].

The full model consists of the following reactions, for a set of *N* expressed gRNAs 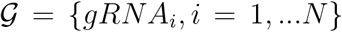 and a set of indices 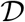 such that every 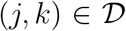 corresponds to a promoter *DNA_jk_* in the network that produces *gRNA_j_* and is repressed by *gRNA_k_* (where *k* is possibly null):

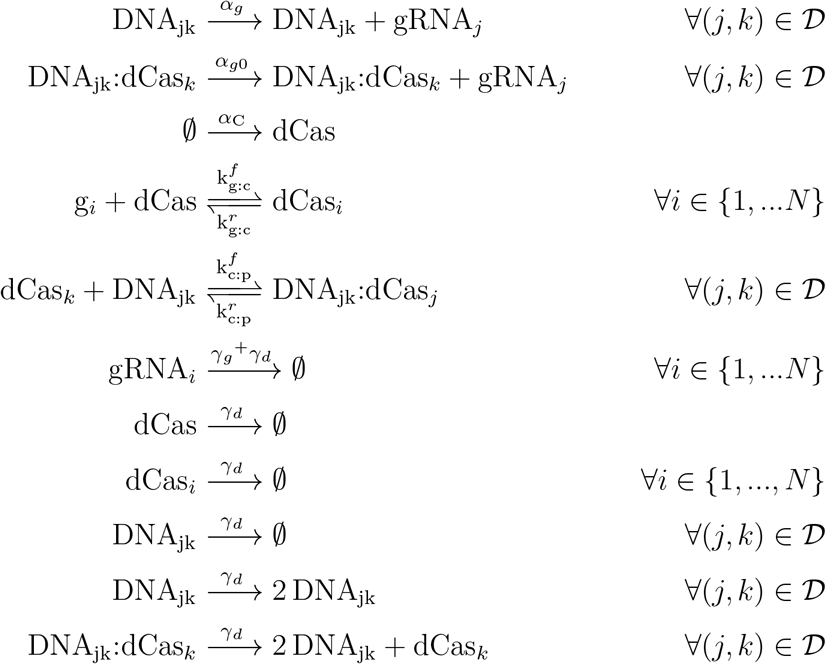

### 2.2. Parameterization of the model

A set of typical parameters used in simulations below is given in Table 1. DNA concentrations will be assumed to be 2 nM unless otherwise noted, which roughly corresponds to the concentration of a genomically-integrated CRISPRi system in actively growing *E. coli* cells.

It is worth mentioning that there is still a great deal of uncertainty around the kinetics of dCas binding. The rates given above for binding and unbinding of gRNAs to dCas were taken from measurements of dCas/gRNA association rates *in vitro* [16]. It is unclear how closely this estimate follows *in vivo* kinetics. For example, the same authors show that the addition of total human lung RNA to an *in vitro* dCas:gRNA assembly reaction slowed dCas:gRNA binding by at least an order of magnitude. Therefore, the estimate in Table 1 may be an optimistic one.

Rates of association between gRNA-loaded dCas and its DNA targets have been more widely studied, but there the literature is still conflicted on their actual values. For example, [19] reports an unbinding rate of dCas from DNA of about 1/(6.5 min) in a radiolabeled pulse chase assay, but that is incompatible with the observation of [20] that dCas dissociation *in vivo* is driven by cell division, or with the real-time, single-molecule measurements of [18], who could not observe sufficient unbinding events over several hours to estimate an unbinding rate for matched gRNAs (setting an upper bound on unbinding time on the order of hours). The parameters for dCas:DNA interactions chosen in Table 1 use binding rates from [16], and reflect the canonical understanding in the field that, for all practical purposes, dCas does not unbind from DNA.

**Table 1:**
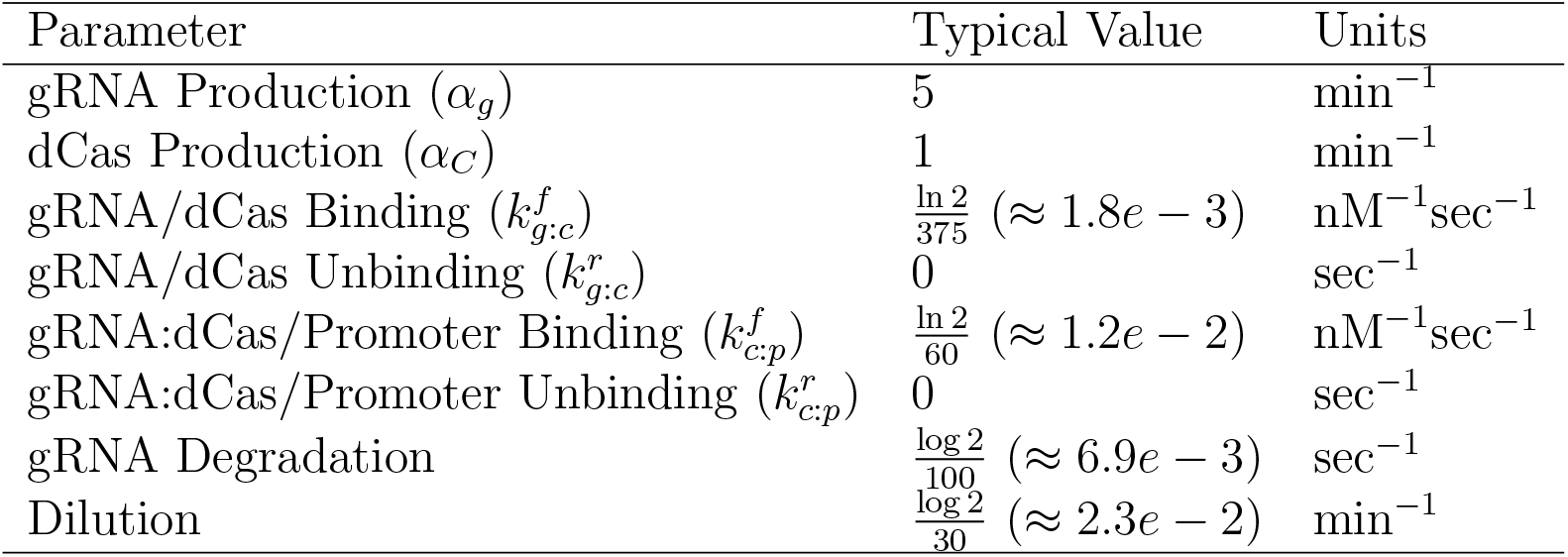
Typical parameter values used for simulating CRISPRi circuits. Parameters are estimated from the literature, except where noted in the text

## 3. Modeling Results

The full CRISPRi model predicts that a variety of dynamic circuits can be constructed from CRISPRi, including a repressilator (though not for all initial conditions; see Section 3.2), a toggle switch, a pulse generating type I incoherent feed forward loop (IFFL), and multiple IFFLs independently driven by a 5-node oscillator (Figure 2). However, these circuits do not operate well under all possible (or even all “reasonable”) parameter values.

**Figure 2:**
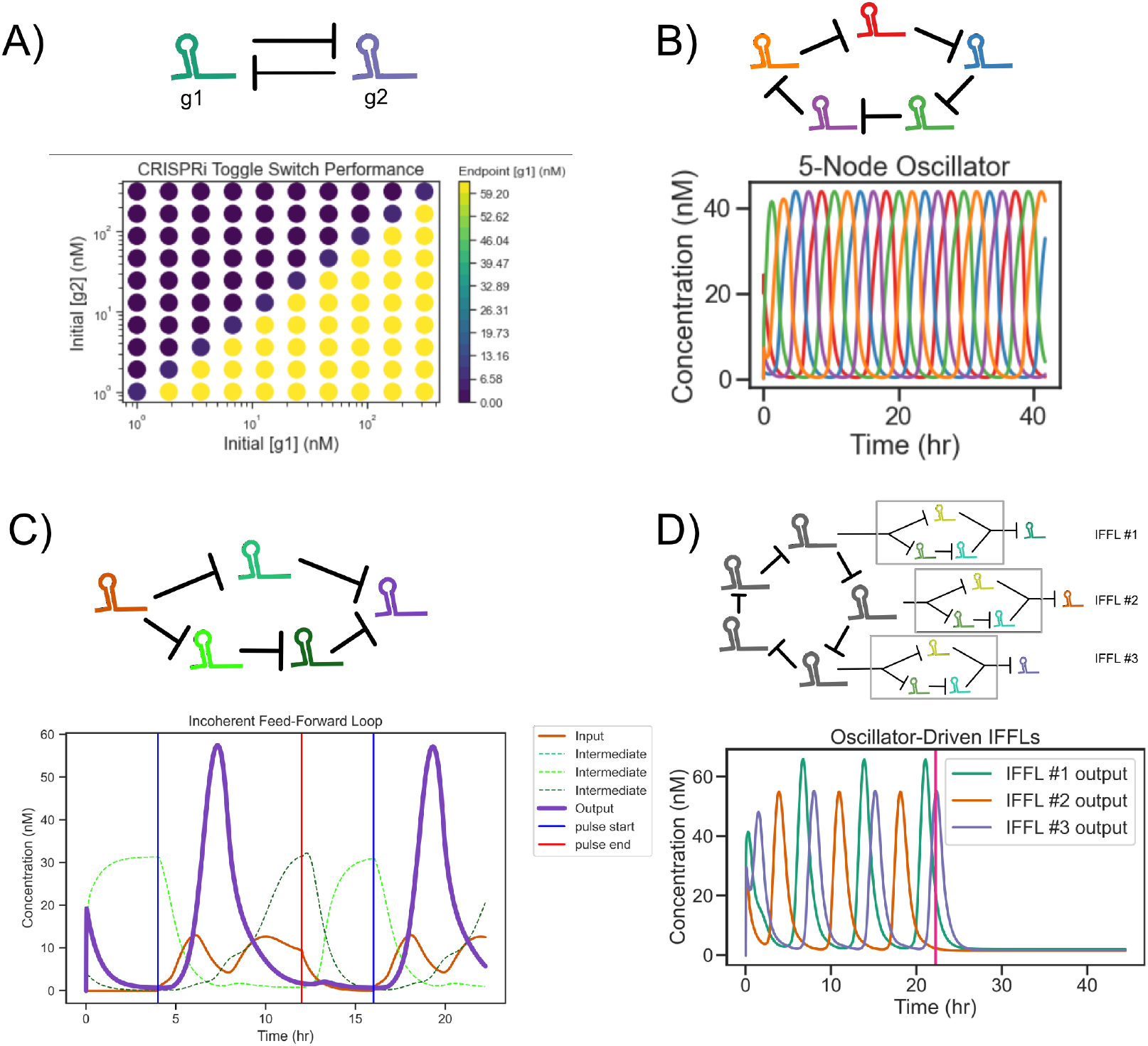
Four examples of dynamic circuits made from CRISPRi components, simulated with the full CRISPRi model. Nodes in circuit diagrams represent gRNA expression units; blunted arrows represent dCas-mediated repression. (**a**) Steady states of a toggle switch for a variety of initial conditions of gRNA concentration. (**b**) A 5-node oscillator. (**c**) A type-I IFFL (pulse generator). The purple trace tracks the IFFL output. Vertical blue and red lines mark activation and return to baseline of the input gRNA promoter, respectively. (**d**) Outputs of three IFFLs driven by a 5-node oscillator. When the node of the oscillator driving the IFFLs is removed (vertical red line), pulses cease. Note that not all nodes of the oscillator have corresponding visible IFFL outputs, and that the peak heights of the three IFFLs are not symmetric.

To better understand the parameter requirements of CRISPRi circuits, and the kinds of engineering we might perform to improve CRISPRi circuit robustness, we will start by using a simple approximation of the CRISPRi model before moving to simulations using a more complete model under varying parameters.

### 3.1. An approximation

Can we understand CRISPRi dynamics in rational, analytical terms? Should we expect an oscillator made from CRISPRi components to actually oscillate? A toggle switch? An IFFL? According to traditional genetic circuit analysis, the toggle switch [8] and repressilators [1] rely on cooperative binding. There is no obvious “cooperative” mechanism in the CRISPRi model, so we might wonder whether we should expect these circuits to function at all.

Unfortunately, the full CRISPRi model outlined in Section 2.2 is not particularly amenable to analysis—even the steady state binding between a single gRNA, dCas, and the gRNA’s target is barely analytically tractable without making an unrealistic quasi-steady state assumption (finding it requires the roots of a rather messy fourth-order polynomial). To attempt to make some headway, we split the model into those parts making up an “idealized,” easy-to-analyze CRISPRi process (informally, “first-order” considerations, though this should not be taken to imply linearity) and kinetic considerations that make CRISPRi difficult to analyze (“second-order” considerations).

We propose the following assumptions for a first-order CRISPRi model: dCas is always present in abundance relative to both DNA targets and gRNAs; binding between gRNAs, dCas, and DNA is instantaneous; and binding of dCas to DNA targets is irreversible. Under these assumptions, the binding of gRNA to dCas to target DNA reduces to a simple “linear” model—with increasing concentrations of gRNA, dCas binds 1:1 with DNA until the DNA is completely saturated. This simplification obviously neglects some important features of CRISPRi (binding kinetics and loading effects on dCas, to name two), but it can still provide insights into how CRISPRi circuits work (or don’t).

Consider a CRISPRi toggle switch consisting of two gRNAs repressing each other. The first-order model of the CRISPRi toggle switch can be modeled with just two differential equations:

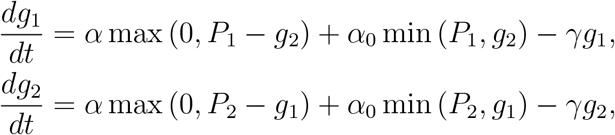

Here, *g*_1_ and *g*_2_ are concentrations of two mutually-repressing gRNAs, *P*_1_ and *P*_2_ are total concentrations of promoters for those guides, *α* is the production rate of gRNA from an unbound promoter, *α*_0_ is the production rate of gRNA from a bound promoter (leak), and *γ* is the division rate of the cell (dilution).

Under what conditions does this system admit two stable steady states? To answer this, we should consider the intermediate steady state of the system, far from the bounds set by 0, *P*_1_, and *P*_2_. In general, toggle-switch-like circuits undergo a supercritical pitchfork bifurcation at this point. When it is stable, the system admits only one state (Figure 3A), but when it is unstable, the system will have two steady states (the “togglable” steady states) (Figure 3B). In particular, the middle steady state will be unstable (and the toggle switch will correctly “toggle”) if and only if that system has a single non-trivial steady state that is unstable. This corresponds to the case where at least one of the eigenvalues of the Jacobian of the system has positive real part. To find when this is true, we note that far from any saturating bounds (where we are likely to find the central steady state), the system reduces to

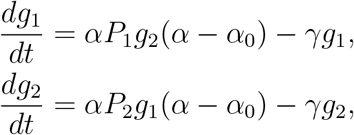

whose Jacobian has eigenvalues – (*α* – *α*_0_) – *γ* and (*α* – *α*_0_) - *γ*. The second eigenvalue always has negative real part. The first eigenvalue has positive real part (and the system “toggles”) when *α* – *α*_0_ > *γ*. In short, a toggle should function as long as the difference between production rates of bound and unbound promoters is sufficiently large relative to dilution.

**Figure 3:**
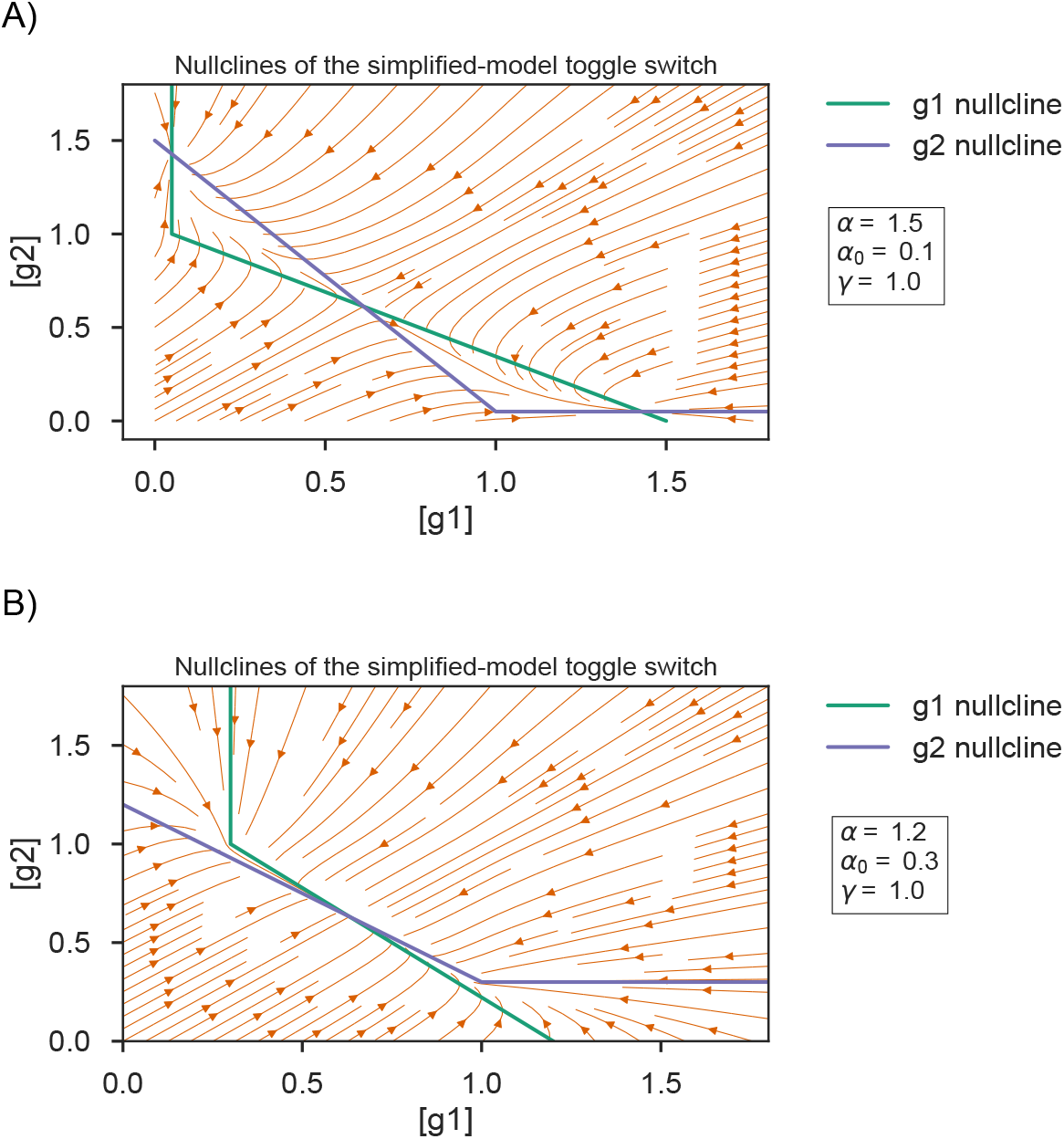
Flow fields for the first-order approximation model of a CRISPRi toggle switch. Depending on the parameters chosen, the toggle could (**a**) admit two stable steady states separated by an unstable steady state or (**b**) admit a single stable steady state.

We can apply a similar analysis to a three-node CRISPRi repressilator, which is an oscillator consisting of an odd number of guide RNAs in a circular circuit topology, each gRNA repressing the next in the cycle (similar to Figure 2B). Bounded dynamical systems with repressilator-like architecture typically have a single non-trivial steady state. As with bistability in the toggle switch, oscillations can occur only when that central steady state is unstable–unless, as we will see, there is also an unstable limit cycle enclosing the steady state. In principle, an unstable central steady state is not sufficient to guarantee oscillations; in practice, molecular species concentrations are bounded by dilution, and there are no other possible stable steady states to the repressilator system, which leaves little room for non-oscillitory (or chaotic) behavior.

The Jacobian for a three-node CRISPRi repressilator has eigenvalues 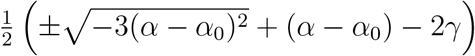 and – (*α* – *α*_0_) – *γ*. The last eigen-value always has negative real part. The first pair of eigenvalues each have positive real part (and the system oscillates) when 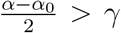. As with the toggle switch, the difference between production rates of bound and unbound promoters must be sufficiently great for the CRISPRi repressilator to oscillate.

We will soon see that this simplified model is a poor predictor of exactly what behavior a CRISPRi circuit with particular parameters will or will not exhibit. This is beside the point; what we learn from the simplified CRISPRi model is that cooperativity is *not* necessary for either a toggle switch or a repressilator, despite classical understanding in the literature [8, 28]. Cooperativity, it seems, is necessary only when genes are expected to bind in a Hill-like fashion. Perfectly linear binding with sharp saturation, as we should expect in CRISPRi, is another perfectly viable path to useful instability.

Now that we have some theoretical justification for believing that a CRISPRi toggle switch or repressilator should be possible to build, we will explore what conditions allow those circuits to function under the full CRISPRi model.

### 3.2. Repressilators can be made with CRISPRi, but they can display strong initial condition dependence

Our full model predicts that the 3-node CRISPRi repressilator, or CRISPRlator, can oscillate slowly, with a period of about seven hours (about three times as long as the original protein-based repressilator [1]). The CRISPRlator’s speed is set by the time scale with which dCas:gRNA complex is removed from the system, which is set by dilution rate.

Interestingly, even under parameterizations that allow oscillations, the CRISPRlator does not oscillate for all initial conditions (Figure 4). It is possible for the 3-node CRISPRlator to possess both a stable limit cycle and an unstable limit cycle inside the stable limit cycle which screens off trajectories with insufficient differences in concentrations of different gRNAs. These latter trajectories, which fall inside the unstable limit cycle, spiral to a stable steady state.

**Figure 4:**
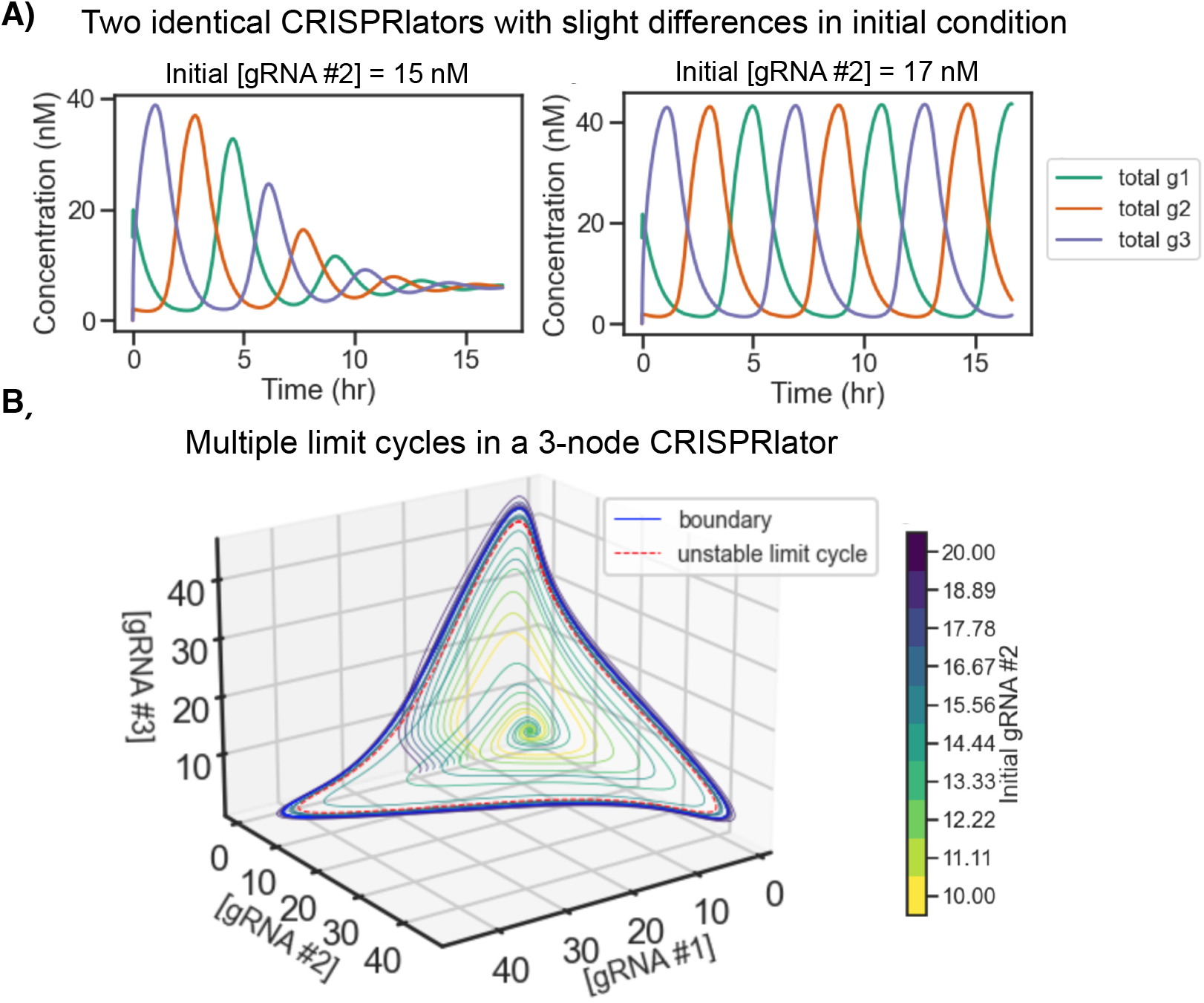
**(a)**The same CRISPRlator (with the same rate parameters) oscillates for some initial conditions but not for others. In both cases, dCas begins at 43 nM and all other non-DNA species other than gRNA #2 start at 0 nM. **(b)** Trajectories for a variety of initial conditions reveal multiple nested limit cycles in the 3-node CRISPRlator under some parameterizations. In this example, an unstable limit cycle (dotted red loop) screens trajectories with too little total gRNA from a stable limit cycle (solid blue loop).

Note that this behavior differs from the behavior of the classic repressilator, which (in its reduced three-species form) can only have up to a single, stable limit cycle [29, 30]. Also note that the existence of multiple limit cycles in the CRISPRlator implies that linearization characterization of the steady state of a CRISPRlator is insufficient to determine whether the CRISPRlator will actually oscillate. If the CRISPRlator’s steady state is unstable, then it will oscillate; but if the steady state is *stable*, then it could either have a pair of stable/unstable limit cycles (in which case it can oscillate), or no limit cycles (in which case it will not). Accordingly, when we computationally screen for oscillations in various CRISPRlators, as in Figure 6, we do so using numeric simulation and heuristic oscillation detection rather than with stability analysis.

### 3.3. The CRISPRlator is fragile

Unfortunately, the CRISPRi repressilator is fragile, and appears to sit close to a bifurcation in parameter space. It is, for example, fairly sensitive to dCas production rate, and ceases to oscillate with less than about 10% or more than about 150% of the default dCas production rate in Table 1. This indicates that dCas production levels may have to be fine-tuned specifically for any particular CRISPRi circuit. Luckily, this is a relatively easy parameter to tune in a real cell.

The repressilator is also not generally robust against transcriptional leak. The first-order model predicts that an increase in leak should stabilize the system towards a steady state, eventually driving it to equilibrium with no oscillations. This is reflected in the full model, and not only for the CRISPRlator—addition of as little as 1% leak destroys oscillations in all but one of the simulations shown in Figure 2 (the exception is the CRISPRlator-driven triple IFFL of Figure 2D, which breaks between 2 and 4% leak).

Empirical evidence is consistent with the idea that the CRISPRlator is buildable but fragile. At least two different dCas9-based CRISPRlator systems have been built to date, and both display oscillations with distinctly irregular amplitude and frequency (Figure 5A-B) [25, 26]. We used stochastic simulation to more fairly compare our CRISPRlator model to experimental data, which required some careful reworking of our model’s handling of plasmid replication (see [27] for details). Although our model predicts that CRISPRlators *can* achieve smooth, regular oscillations even under stochastic kinetics, with tuned dCas production rate, we can reproduce the empirically-observed balance between consistent oscillation and amplitude/wavelength variance (Figure 5 C-D). At high dCas production rates, the CRISPRlator oscillates relatively cleanly; at low dCas production rates, the CRISPRlator breaks down into apparently random bursting; at moderate dCas production rates, oscillations occur consistently but with irregular profiles, as appears to happen in lab-built CRISPRlators.

**Figure 5:**
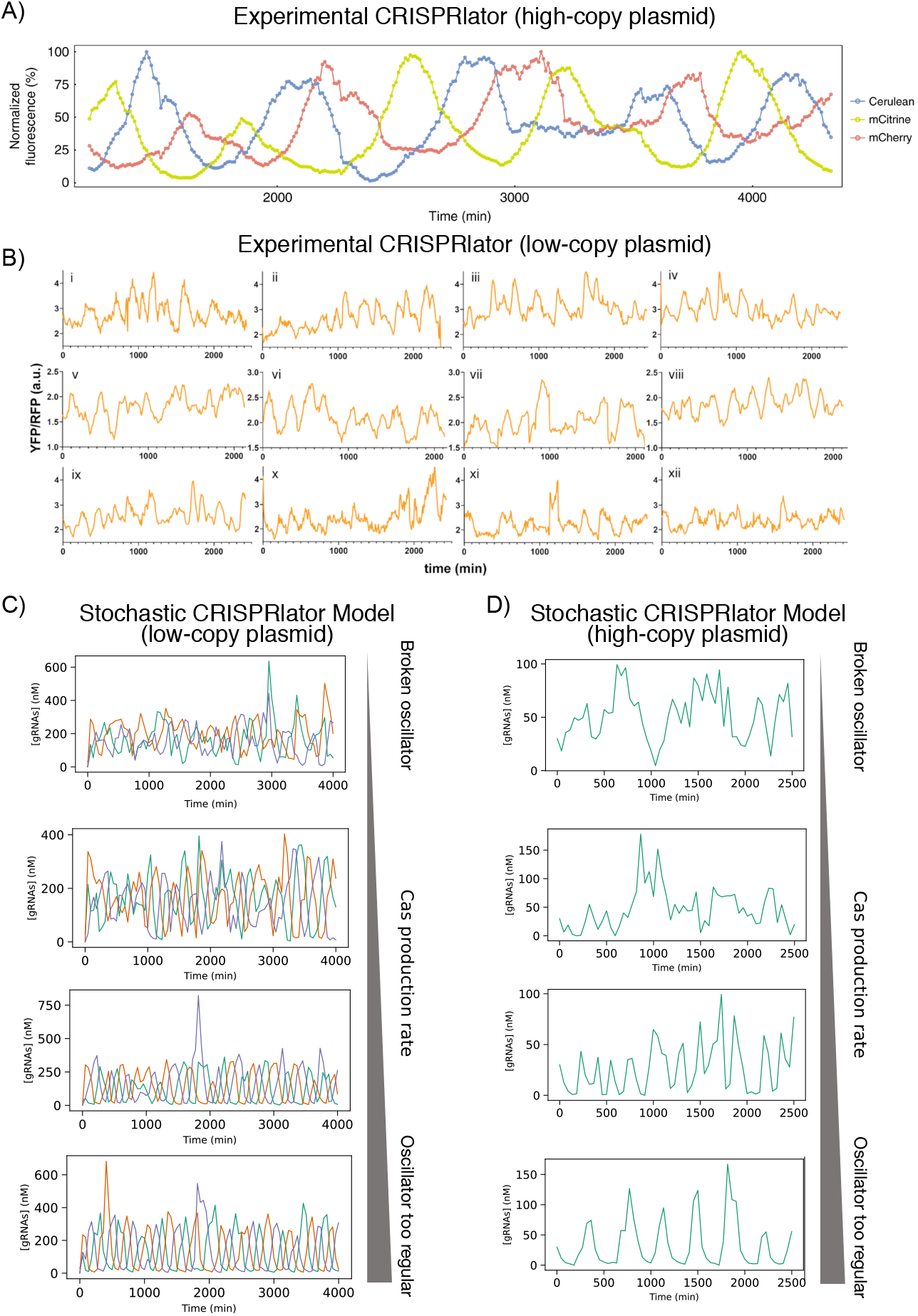
**(a)** Example real-world CRISPRlator oscillations from a high-copy plasmid (reproduced from [25]). **(b)** Example real-world CRISPRlator oscillations from a low-copy plasmid (reproduced from [26]). **(c-d)** Stochastic simulation of our model CRISPRlator. Lowering dCas production rate smoothly increases noise in observed oscillations, and can be roughly matched to the experimental data shown in (a) and (b), respectively.

Notably, our simulations suggest that a CRISPRlator should be very difficult to build if there exists any transcriptional leak from dCas-bound promoters. That a CRISPRlator has, in fact, been built successfully suggests that CRISPRi might, in fact, have extremely low leak in the true sense of allowing transcription while the repressor is bound. Repression with dCas has been reported in cell-free extract with fold-repression between 7 and 100 [18], and *in vivo* with similar repression strengths [13, 15], which puts the best CRISPRi repression in a leak range that should not be likely to allow a repressilator to work. Our model of the CRISPRlator suggests that the “leak” observed in these experiments may be a function of *slow binding*, not failure of a bound repressor—that is, observed “leak” occurs because a promoter is not bound some of the time, not because a promoter is bound to dCas and produces transcripts anyway.

### 3.4. Well-tuned production and degradation of dCas can offset leak-based circuit fragility

There are a few different knobs we can turn to make the CRISPRi repressilator more robust to transcriptional leakiness. We can decrease the rate of production of either dCas or gRNAs; we can speed the binding between dCas:gRNA complexes and DNA (in contrast, speeding binding between dCas and gRNAs appears to have little effect); we can add active degradation of dCas (also increasing the *speed* of the oscillator considerably); and we can grow the repressilator from three nodes to five nodes.

As an example, let us consider the interaction between dCas degradation rate and leak rate. Figure 6 shows the performance of simulations of several circuits as a function of the degradation rate and leak rate parameters, with other parameters as shown in Table 1. We can think of these charts as a sort of two-dimensional “specification sheet” for dCas to allow different circuits to function properly. For any particular leak rate, the circuit will either oscillate (Figure 6A, B, D, E, and F) or toggle (Figure 6C) only when dCas is degraded at a rate within a proper range. Too much dCas degradation will destroy all examined circuits, and a minimum amount of degradation is required for some. Notably, the “correct” dCas degradation specification is different for different circuits (compare Figures 6A, 6B, and 6C), dCas expression rates (Figures 6D and E), and background activity (different numbers of oscillators in Figure 6F). Interestingly, the toggle switch has similar parameter requirements on these two axes, suggesting that there may be some requirements shared by some interesting class of dynamic CRISPRi circuits.

**Figure 6:**
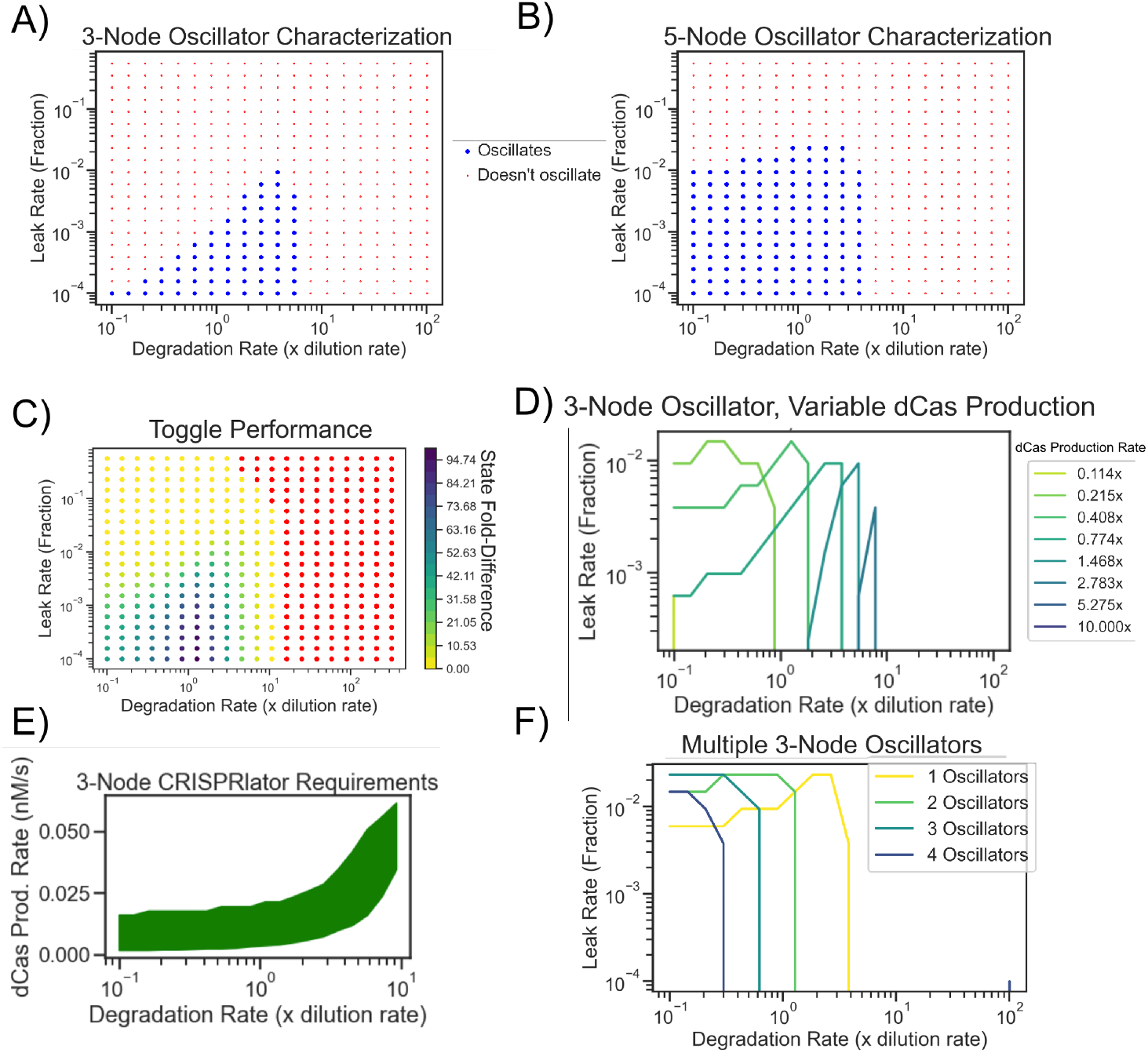
Acceptable degradation rates vary by circuit architecture and parameterizations. Leak is given in units of the rate of gRNA production from an unbound promoter. Degradation rate is in units of cell division rate. Colors at each point in parameter space indicate whether the circuit oscillates (blue) or not (red) for those given parameters. Results are given for **a)** a 3-node CRISPRlator, **b** a 5-node oscillator, and **c)** a CRISPRi toggle switch. In **c**, Colors indicate the separation distance between the two gRNAs at steady state; red indicates no detectable bistability. **d)** Changes in degradation requirements for a 3-node CRISPRlator with different levels of dCas expression. Areas under each curve represent parameters for which the circuit oscillates. Production rates are given in units of min^-1^ (the “default” dCas speed used in (textbfa)). **e)** Changes in dCas expression requirements for the 3-node CRISPRlator with different rates of dCas9 degradation. The shaded region represents production rates that admit oscillations. Oscillations could not be recovered with any higher degradation rate. Transcriptional leak is set to 0. **f)** Parameters for which different numbers of independent 3-node CRISPRlators oscillate while operating in the same cell. Larger number of oscillators become steadily less tolerant of both degradation and leak.

We can produce a similar “specification sheet” for dCas degradation and leak for a 5-node CRISPRi repressilator, as shown in Figure 6B. The 5-node repressilator is more robust than the 3-node oscillator. Indeed, the 5-node repressilator can operate with as much as 10% leak or as little as no dCas degradation at all. In the case of CRISPRi repressilators, bigger is not only better, but potentially easier.

On the other hand, the parameter requirements of the toggle switch appear to be quite similar to those of the 3-node repressilator (Figure 6C). Admittedly, the toggle switch and 3-node repressilator have very similar architecture, but the fact that both circuits require similar degradation rates and minimum promoter leak suggests that the regime of functional repressilators may have not-yet-understood underlying properties that are broadly useful for constructing CRISPRi circuits.

There are more than two tunable knobs in the CRISPRi system. One that we have already seen to be important is the *production* rate of dCas. Figure 6D shows how the target parameter set changes with different levels of dCas production. The good news is that with low enough dCas expression there is no need for dCas degradation (though with dCas steady state levels that low, stochastic fluctuations become a more serious problem). The bad news here is that at least one *engineerable* but not *readily tunable* parameter of CRISPRi (namely, dCas degradation rate) has acceptable value ranges that do not overlap for some choices of dCas production rate. This should not be too much of a problem for making a single repressilator, but it does complicate the design and integration of multiple CRISPRi circuits in the same cell. For example, Figure 6E shows the expected effect of expressing two identical CRISPRi repressilators in parallel with no directly cross-interacting nodes. The increased load on dCas drops the effective steady-state concentration of dCas as perceived by each individual oscillator, which has a similar effect as dropping dCas production rate. Namely, this shifts the required rate of dCas degradation. A repressilator that works on its own can be expected to fail when a second repressilator is added, unless dCas’s degradation rate is exquisitely well-tuned. More generally, it seems likely that different circuits may require dCas variants with different degradation rates.

Reciprocally, we could tune degradation rate to compensate for changes in other parameters. The CRISPRlator requires dCas to be produced at a tuned rate—too much or too little dCas production will destroy the circuit’s function. Changing the rate at which dCas is degraded *also* changes those dCas production rate requirements, potentially allowing degradation to compensate for any lack of control over dCas concentration (Figure 6D).

Finally, we can consider the robustness of the CRISPRlator to differences in the strengths of the promoters driving gRNA production. In general, repressilators require nodes with roughly similar repression strengths. The more sensitive the CRISPRlator to gRNA promoter strengths, the more difficult a CRISPRlator will be to engineer. Fortunately, as shown in Figure 7, the 5-node CRISPRlator is robust to (most) changes of at least 10-fold in two adjacent gRNA.

**Figure 7:**
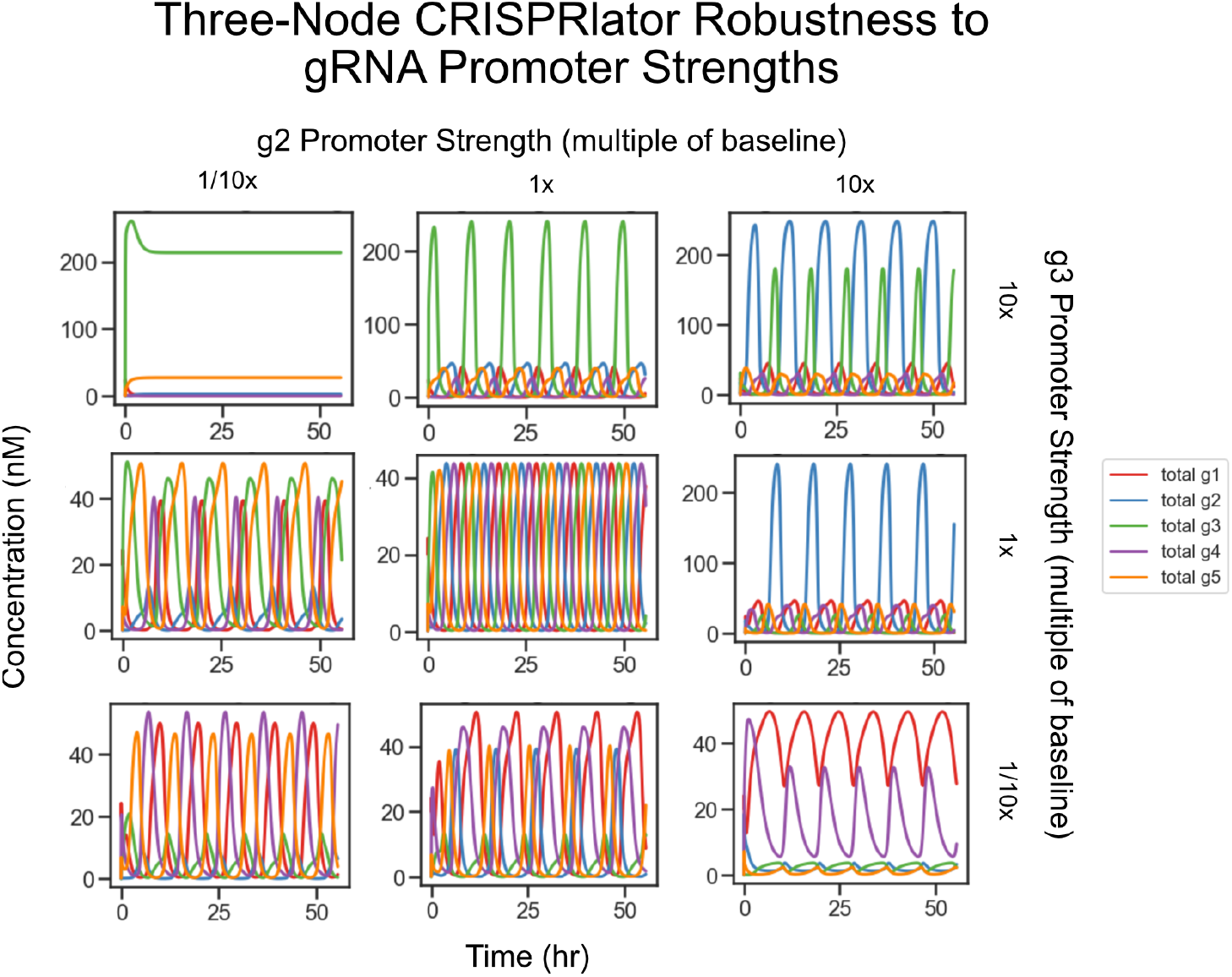
The 5-node CRISPRlator functions under many possible changes to the strengths of two adjacent gRNA promoters. Other parameters are as in Figure 2b. Each panel shows simulation results with one combination of promoter strengths.

## 4. Engineering Requirements of CRISPRi

Although we do not yet have a complete set of guidelines for engineering arbitrary CRISPRi systems or combinations of systems, our simulations provide a few key lessons:

- Active degradation of dCas can improve circuit performance, and may be necessary.
- Almost any amount of transcriptional leak can be quite destructive to CRISPRi circuits.
- The speed at which dCas:gRNA complexes bind to DNA is important for circuit function, and implementing CRISPRi circuits *in vivo* may require increasing this binding rate.

Each of these lessons is accompanied by an engineering requirement. Some possible strategies for fulfilling these requirements are outlined below.

### 4.1. Degradation of dCas

*E. coli* can actively degrade proteins using the ClpXP system, which uses the proteases ClpXP and ClpXA to selectively degrade proteins bearing ssrA, a small C-terminal peptide tag. The native ClpXP system degrades at least 400 times faster than the rate of cell dilution, which is far too fast for any of the circuits outlined in Section 3 [31]. However, targeted mutations to the ssrA tag have been used to tune degradation rates to anywhere between 2 and 100 times the rate of dilution, which falls solidly into the target degradation range outlined in Figure 6 [32]. Alternatively, dCas could be degraded using the *mf*-Lon protease system, which is similar to the ClpXP system in function and tunability but is orthogonal to any system in *E. coli* [33, 34].

### 4.2. Improving fold-repression

The CRISPRi repressors reported in the literature typically repress with strengths between 10x and 100x. Moreover, the strongest CRISPRi repressors are typically targeted inside the target promoter, which limits their design space severely. Simulations suggest that leaks reported for elongation-blocking CRISPRi will break toggle switch and oscillator circuits unless degradation rates are extremely well-tuned. Engineering high fold-change in a promoter may be challenging, but several simple strategies are possible for improving repressor performance.

However, any “transcriptional leak” of the kind observed in any repression experiment can be explained as a consequence of slow DNA binding or weak binding equilibrium, rather than proper leak from bound promoters; in fact, real-world “leak” is likely to be a combination of the two. Our models predict that CRISPRlators without dCas degradation should only function if proper leak from dCas-bound promoters is extremely low.

Real-world CRISPRlators, though not particularly robust, do oscillate, suggesting that a consequence of our model would be that proper leak in CRISPRi systems is much smaller than leak from un-bound targets. If true, this suggests that efforts to reduce CRISPRi leak should be targeted at improving binding speed, rather than binding strength.

### 4.3. Faster dCas:DNA binding

As previously mentioned, the results shown in this report assume “best-case” binding rate of dCas:gRNA complexes to their target DNAs, with binding rates taken from *in vitro* association rate measurement. In live *E. coli*, dCas binding to DNA is much slower, with a single dCas complex estimated to require a few hours to bind to its target [20]. A likely explanation for Cas9’s slow binding time is that dCas spends most of its time transiently bound to off-target PAM sites in the genome. The dCas protein can temporarily bind to any double-stranded DNA site with a correct PAM sequence. When it does, it briefly opens the DNA helix to “check” whether its associated guide RNA matches the sequence immediately adjacent to the PAM. Each non-target PAM present in the cell slows the rate of correct binding by acting as a low-affinity “decoy” binding site, which slows binding to the target promoter. The *S. pyogenes* Cas9 (by far the most widely-used Cas variant, and the one used exclusively in this report) uses the PAM “NGG”, which can be expected to appear roughly 750,000 times in the (approximately diploid) genome of a growing *E. coli* cell. This represents substantial barrier to correct target identification.

There are several possible solutions to the problem of slow dCas:DNA binding *in vivo*. The most straightforward, at least conceptually, would be to move out of cells entirely and construct circuits exclusively in TX-TL or another cell-free system. Since cell-free systems do not have DNA replication or dilution to remove dCas from DNA, this would have to be done either in a microfluidic device capable of manually diluting a running reaction (see [35]) or using degradation-tagged dCas proteins.

Another simple way to speed up dCas:DNA binding rates would be to simply increase the concentration of either dCas (by increasing the baseline production of dCas) or target (by either genomically integrating multiple copies of the CRISPRi circuit or by expressing the circuit off of a plasmid). Simulations so far suggest that, all other things being equal, increasing the rate of dCas production has the effect of narrowing the window of acceptable dCas degradation rates. More simulation will be required to determine the feasibility of either of these two interventions.

Another possible solution would be to use a dCas (or another programmable binding protein) with a different, more complex PAM sequence. For example, the *Treponema denticola* Cas9 (TD-Cas) uses the PAM NAAAAC. After accounting for nonspecific PAM recognition, TD-Cas PAM sites ought to occur between 64 and 1024 times less frequently than *S. pyogenes* Cas PAM sites, with a corresponding increase in binding rate [12]. If the *other* kinetics of TD-dCas are similar to those of *S. pyogenes* dCas, then we should expect TD-dCas binding to DNA to be only perhaps twice as slow as *in vitro* dCas, which should be quite manageable.

## 5. Conclusions

CRISPRi remains an intriguing technology for scaling up genetic regulatory networks. However, building functional CRISPRi circuits is not as simple as sketching a repression net and targeting gRNAs against each other accordingly. In particular, ODE simulations of CRISPRi reveal specific functional requirements regarding dCas9 protein regulation and repressor characteristics. Some degradation of dCas may be required, but not *too* much; some leak from dCas-repressed promoters is acceptable, but not *too* much. Long timescales of DNA binding make CRISPRi construction more difficult, but not fatally so. Using these and other insights, it should be an achievable goal to build and express CRISPRi circuits, which would constitute an important milestone toward engineering cells with complex programmable behavior.

## 6. Acknowledgments

This work was supported by the Human Frontiers Science Program, by National Science Foundation award number 1317694, and by the Institute for Collaborative Biotechnologies through contract W911NF-19-D-0001 from the U.S. Army Research Office.

